# A systematic review and meta-analysis of the human blood index of the major African malaria vectors

**DOI:** 10.1101/424093

**Authors:** James Orsborne, Luis Furuya-Kanamori, Claire L. Jeffries, Mojca Kristan, Abdul Rahim Mohammed, Yaw A. Afrane, Kathleen O’Reilly, Eduardo Massad, Chris Drakeley, Thomas Walker, Laith Yakob

**Author notes:** LY contact: Department of Disease Control, London School of Hygiene & Tropical Medicine, Keppel Street, London, WC1E 7HT.

## Abstract

**BACKGROUND:** The proportion of mosquito blood-meals that are of human origin, referred to as the ‘human blood index’ or HBI, is a key determinant of malaria transmission. We conducted a systematic review of the HBI for the major African malaria vectors.

**RESULTS:** Evidence is presented for higher HBI among *Anopheles gambiae* (M/S forms and *An. coluzzii*/*An. gambiae s.s.* are not distinguished for most studies and therefore combined) as well as *An. funestus* when compared with *An. arabiensis* (prevalence odds ratio adjusted for collection location [i.e. indoor or outdoor]: 1.62; 95%CI 1.09-2.42; 1.84; 95%CI 1.35-2.52, respectively). This finding is keeping with the entomological literature which describes *An. arabiensis* to be more zoophagic than the other major African vectors. However, analysis also revealed that HBI was more associated with location of mosquito captures (R^2^=0.29) than with mosquito (sibling) species (R^2^=0.11).

**CONCLUSIONS:** Our findings call into question the appropriateness of current methods of assessing host preferences among disease vectors and have important implications for strategizing vector control.

## Background

Malaria is transmitted through mosquito bites, making the vectors’ choice of which blood-host species to bite a central component of malaria epidemiology and ecology. In Africa, the majority of infections are transmitted by *Anopheles gambiae* sensu stricto, *An. coluzzii*, *An. funestus* and *An. arabiensis*. Conventional wisdom indicates that the first three vectors are anthropophagic while the latter sibling species is more zoophagic. Levels of anthropophagy/zoophagy are typically assessed using PCR to identify the host species from blood-meals in field-caught mosquitoes, and are then quantified according to the human blood index (HBI), defined as the proportion of blood-meals that are of human origin [1]. Because two mosquito bites on a human are required to complete the malaria parasite’s life-cycle, HBI has an inflated impact on metrics of transmission such as the basic reproduction number, the vectorial capacity and the critical density of mosquitoes for sustained transmission [2].

However, the HBI should not be perceived to have a singular, fixed value; all major African malaria vectors have demonstrable plasticity in the host species that they bite [3–5]. It has long been recognised that the same mosquito population will often adjust its biting towards a more locally available host species [1, 6]. This has important implications for malaria control policy. For example, recent studies have observed that increased outdoor biting followed the distribution of insecticide treated bednets [7]. In such circumstances, vector control tools that operate effectively outdoors become a critical component for eliminating local malaria transmission. Unfortunately, the huge malaria burden reduction achieved in the years since 2000 has relied disproportionately on control tools operating indoors [8], and there are limited effective malaria-vector control options for outdoor use.

One technology that shows promise for targeting mosquitoes regardless of whether they bite indoors or outdoors involves the use of systemic insecticides – chemicals applied directly to blood-hosts to kill mosquitoes that take a blood meal. This technology arose from the observation that mosquito mortality was increased following the consumption of sugar-meals [9] or blood-meals [10] containing ivermectin - a drug used for onchocerciasis control. Drugs approved for veterinary use, such as fipronil, have subsequently been demonstrated to have similar impact when livestock are dosed orally, or when the chemical is applied topically [11]. More recently, systemic insecticides have had durations of their efficacy extended through dosing with higher concentrations [12], combined dosing with adjuvants [13], and with use of sustained-release devices [14]. The stage is set for progress in development and evaluation of ivermectin for vector control [15]. Therefore, we argue that it has never been more important to understand the distribution of malaria-vector bites on alternative host species. Here, we systematically reviewed and meta-analysed the current evidence to identify the factors associated with higher HBI in sub-Saharan Africa.

## Methods

Findings from the systematic review were reported following the PRISMA guidelines [16]. The inclusion and exclusion criteria are listed in Table 1 and advanced search terms were developed following initial manual literature searches and a basic PubMed search (Table 2). The purpose of the initial search was to identify keywords and synonyms. The authors agreed on the search terms and inclusion/exclusion criteria before the systematic search was performed. The Ovid database was used to search available MEDLINE and EMBASS literature from inception to February 2018. Books were excluded from all searches as well as articles not written in English. Results were retrieved and collated using Mendeley desktop reference manager.

**Table 1.** Inclusion and exclusion criteria for systematic review.

| Inclusion Criteria | Exclusion Criteria |
| --- | --- |
| Studies which used blood meal analysis (PCR,ELISA or precipitin tests) to report the HBI | Semi field studies, studies using baited traps or choice experiments to investigate host preference |
| Studies performed in sub-Saharan Africa | Entomological studies not specifically reporting the HBI |
| Studies reporting the HBI for individual mosquito species | Studies not reporting total number of mosquitoes caught |
| Reporting HBI for *Anopheles gambiae*, *Anopheles funestus* complex or *Anopheles arabiensis* mosquito species | Data points based on less than 50 blood-fed mosquitoes in total for target species |
| Studies reporting trapping methodology including location of traps (indoors or outdoors) |  |

**Table 2.**
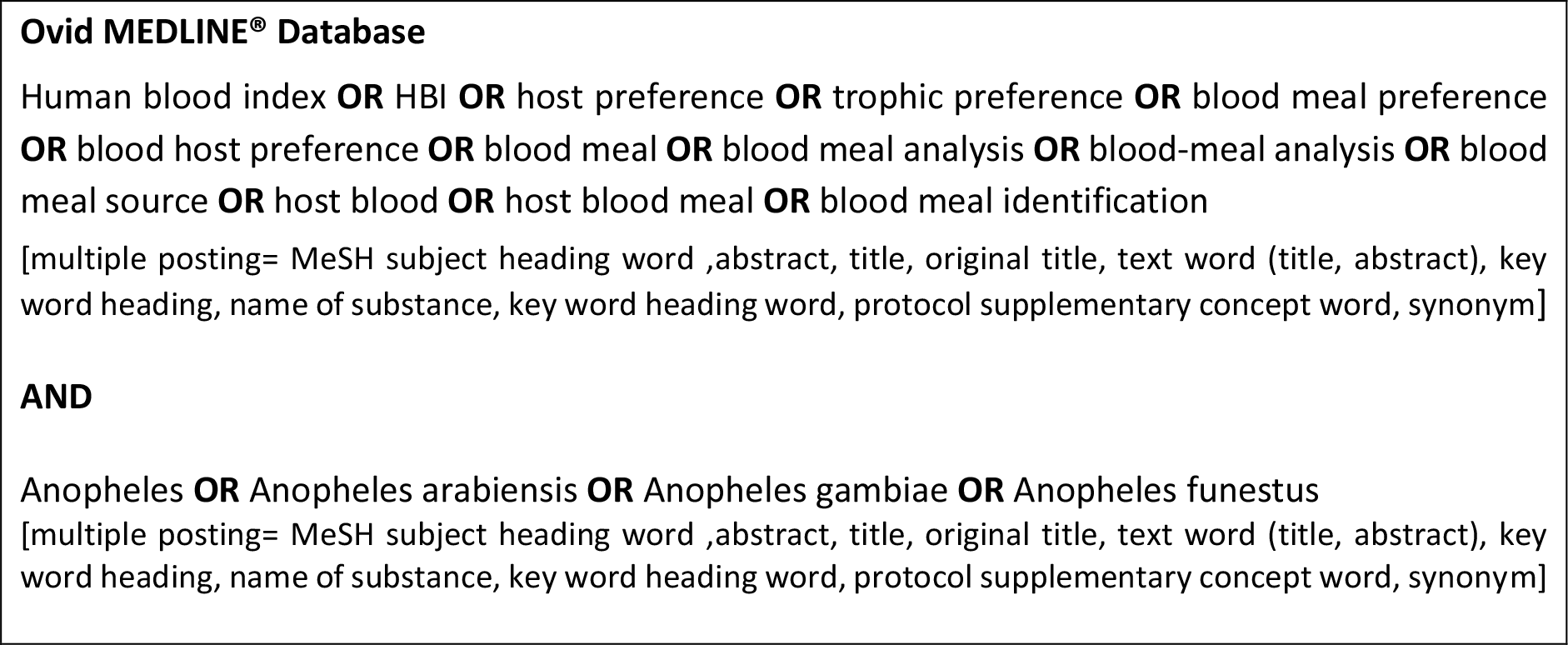
Search strategy for systematic review.

After eliminating duplications, abstracts for all publications retrieved were reviewed for relevance. Full-text reviews were then conducted on all articles to decide on its inclusion in accordance with the pre-specified inclusion and exclusion criteria. If the inclusion criteria were satisfied the estimated Human Blood Index (HBI) reported was retrieved. Other variables that could have a significant effect on the reported HBI were also retrieved. These variables included (sibling) species (complex), trapping location (indoors, outdoors or both), trap type(s) used and total number of mosquitoes collected. The primary effect measure of interest was the HBI.

The double arcsine square root transformed HBI (expressed as a proportion of all blood-meals) was used to stabilize the variance across the studies [17] and then back transformed for ease of interpretation. A linear model was performed on all eligible studies to gain additional insight into the effect of trapping location and *Anopheles* species on the proportion of HBI. The linear model was fit using the HBI (proportion) as the response variable weighted by the inverse of each study’s variance to allow the observations with the least variance to provide the most information to the model, and using robust error variances. All tests were two-tailed and a p-value < 0.05 was deemed statistically significant. Pooled analyses were conducted using MetaXL (version 5.3, EpiGear Int Pty Ltd; Sunrise Beach, Australia) and the regression models were run using Stata MP (version 14, Stata Corp, College Station, TX, USA).

## Results

The search identified 1243 potentially relevant studies. After collating these results and reviewing all abstracts, 662 studies were deemed relevant. All full text articles were retrieved, reviewed for relevance and reviewed against all inclusion and exclusion criteria. Sixty-one studies resulting in 166 data points fulfilled all criteria and where included in the analysis. Reason for exclusion at full text stage included inadequate number (fewer than 50) of mosquitoes collected (n=14) and the use of host-biased trapping methodologies (n=4) (Figure 1).

**Figure 1.**
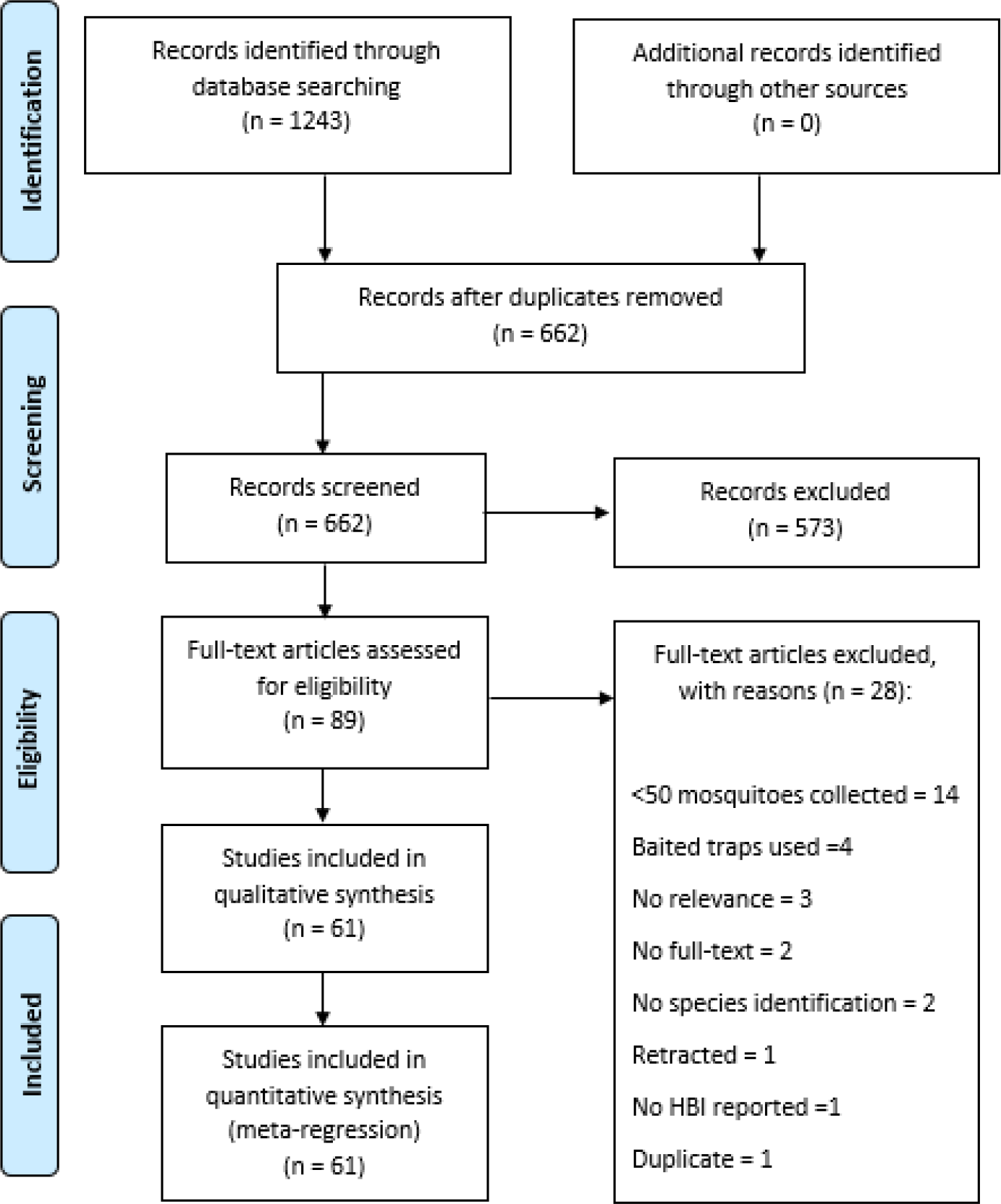
PRISMA flow diagram of search phases with numbers of studies included/excluded at each stage.

Multiple collection methodologies were identified from the eligible studies. The methodology used was governed by the collection location targeted (indoors or outdoors). Indoor collections were the most widely used (n=118) with pyrethroid spray catch (PSC) the most commonly used methodology (n=78). Other collection methods included manual indoor collections (n=20) and the use of CDC light traps within the household (n=10). Outdoor collections represented 27 of the total data points extracted with manual collection of mosquitoes being the most common collection method (n=13). Pit traps (n= 10) and CDC light traps (n=4) were also an effective collection method. Studies collecting from both indoor and outdoor environments consisted of 21 data points. These studies used a variety of different methods; many used a combination of the most effective indoor and outdoor collection methods. CDC light traps were the most common (n= 12) followed by other combinations of indoor and outdoor methods; CDC light trap plus PSC (n=2) and pit traps and manual indoor collections (n=2) (Table 3 and S1). Collection methods had no significant effect (p> 0.05) on the reported HBI when comparing the mean HBI produced by each collection methodology within its respective collection areas (indoor and outdoor) for *An. gambiae, An. arabiensis* and the *An. funestus* species complex (Figure 2). It should be noted that due to the variety of different methods used and therefore sparsity of data for each methodology within the ‘both’ category, a meaningful comparison could not be made.

**Table 3.** Data points extracted from eligible studies for each collection location and trapping methodology. The top three methodologies based on number of data points extracted are displayed here

| Species | Methodology |  |  |  |  |  |  |  |  |  |  |  |
| --- | --- | --- | --- | --- | --- | --- | --- | --- | --- | --- | --- | --- |
|  | Collection Location |  |  | Indoor |  |  | Outdoor |  |  | Both (Indoor + Outdoor) |  |  |
|  | Indoor | Outdoor | Both | Pyrethroid<br>Spray Catch<br>(PSC) | Manual<br>aspiration | CDC<br>light<br>trap | Manual<br>aspiration | CDC<br>Light<br>trap | Pit<br>Traps | CDC light trap +<br>PSC | CDC light<br>trap | Others |
| *Anopheles gambiae* | 37 | 7 | 3 | 22 | 10 | 2 | 7 | 0 | 0 | 1 | 0 | 2 |
| *Anopheles arabiensis* | 50 | 14 | 10 | 39 | 2 | 7 | 1 | 4 | 9 | 0 | 10 | 0 |
| *Anopheles funestus s.l.* | 31 | 6 | 8 | 17 | 8 | 1 | 5 | 0 | 1 | 1 | 2 | 5 |

**Figure 2:**
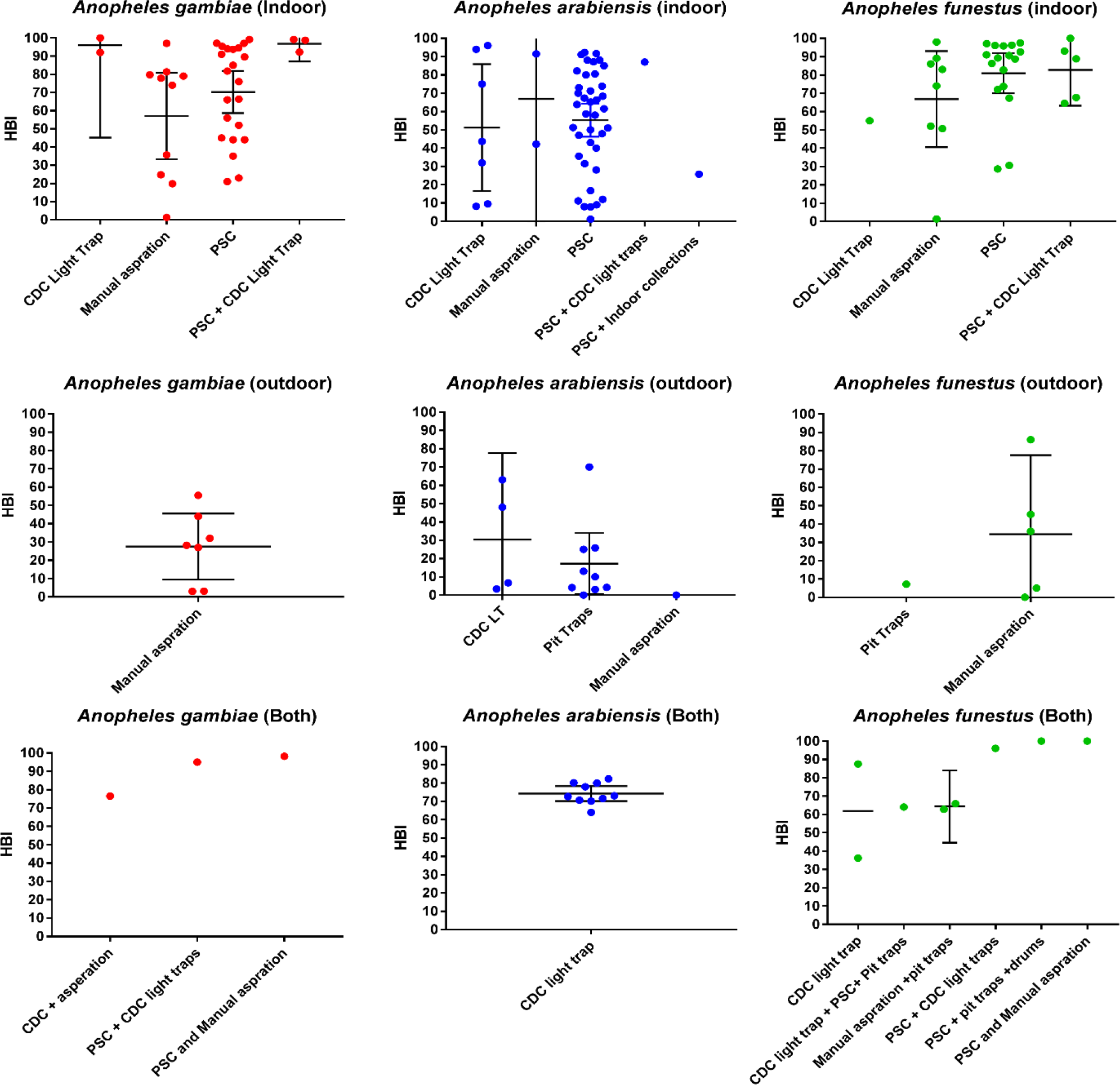
Reported mean HBI (+95% CIs) for individual collection methods when sampling from indoor, outdoor or both (indoor and outdoor) environments for *An. gambiae, An. arabiensis and An. funestus*.

Meta-regression of the data compiled from the 166 data points demonstrated a significantly higher proportion of blood-meals were of human origin (the human blood index, ‘HBI’) among *An*. *funestus* (prevalence odds ratios [POR] of 1.84 (95%CI 1.35-2.52, p<0.001) and *An*. *gambiae* (POR of 1.62, 95%CI 1.09-2.52, p=0.02) compared to *An*. *arabiensis*. The majority of studies including details of *An*. *gambiae* did not specify whether they were M or S forms (or, in more modern nomenclature, *An. coluzzii* or *An. gambiae s.s.*), so these were combined. For all three groups, a significantly higher HBI was found from indoor mosquito collections (POR of 2.74, 95%CI 2.00-3.75, p<0.001) as well as combined indoor and outdoor collections (POR of 4.20, 95%CI 3.13-5.62, p<0.001) versus outdoor only collections. The results also revealed that trapping location (R^^2^^ of 0.29) had a larger impact on the blood-meal host species than mosquito species (or species complex) (R^^2^^ of 0.11) and that this difference was statistically significant (p<0.01 resulting from an F-test comparing both univariate models) (Table 4).

**Table 4.** Predictors of Human Blood Index: univariate and multivariate regression models.

| Predictors | Univariate |  | Multivariate |  |
| --- | --- | --- | --- | --- |
|  | POR (95% CI) | R<sup>2</sup> | POR (95% CI) | R<sup>2</sup> |
| Anopheles species |  | 0.11 |  | 0.40 |
| *An. arabiensis* | 1.00 |  | 1.00 |  |
| *An. gambiae* * | 1.50 (0.95-2.36) |  | 1.62 (1.09-2.42) |  |
| *An. funestus* | 1.95 (1.34-2.85) |  | 1.84 (1.35-2.52) |  |
| Location |  | 0.29 |  |  |
| Outdoor only | 1.00 |  | 1.00 |  |
| Indoor only | 2.83 (2.04-3.93) |  | 2.74 (2.00-3.75) |  |
| Both | 3.98 (3.14-5.06) |  | 4.20 (3.13-5.62) |  |

POR, prevalence odds ratio; \*most studies did not specify M/S form of *An. gambiae* (and predated the renaming of these forms as *An. coluzzii* and *An. gambiae* s.s. respectively).

## Discussion

Control of vector-borne diseases is largely, often entirely, dependent on vector control. For malaria, vector control is achieved primarily through targeting mosquitoes that are host-seeking [8]. The major African malaria vectors, *An. gambiae s.s.*, *An. coluzzii* and *An*. *funestus*, are regularly cited as paragons of anthropophagy, and any non-human biting exhibited by these species has historically been ignored when strategizing control. Here, their biting behaviour was systematically reviewed for the first time and it was demonstrated that the difference in their host choice compared with the zoophilic vector *An*. *arabiensis* was dwarfed by the difference found when comparing indoor with outdoor collections. In other words, *where* the mosquito was collected was substantially and significantly more influential on host choice than *which* mosquito species was collected.

This raises an important question: where should vectors be collected from in order to provide the most useful HBI estimates? Our results indicate that a single HBI for a given location risks presenting quite a biased estimate for local vector biting behaviour. A standardised HBI accounting for both indoor and outdoor behaviours would probably constitute an invalid metric because of the increased difficulty posed by collecting blood-fed mosquitoes outdoors i.e., we lack the tools to estimate indoor versus outdoor mosquito numbers with any confidence. Therefore, our recommendation is to present both estimates for an indoor HBI and an outdoor HBI. Longitudinal assessments initiated before rolling out control tools, and followed up over the time course of the programme would provide a valuable source of information. For example, these would determine the timeframe across which LLIN-derived exophagy [7] occurs, as well as provide unbiased estimates of the magnitude of effect. These entomological data would also be able to inform on whether there is a reversion to behavioural norm after a certain period post-distribution, and the rate at which this occurred.

Better data on this behaviour and its temporality will do much more than inform a fundamental aspect of mosquito ecology: it will have considerable ramifications pertaining to malaria control. For example, if significantly reduced HBI is detected immediately following the distribution of LLINs, this may present an excellent opportunity to synergise bednets with systemic insecticide-treated livestock. Saul [18] described the potential for zooprophylaxis to switch into zoopotentiation if the availability of alternative blood meals increases mosquito survival more than counters the impact of diverting feeds. This risk could be reduced or eliminated with systemic insecticidal dosing that is judiciously timed with LLIN roll-out. Mathematical models already exist for optimal systemic insecticide deployment [19] including its integration with LLINs [20]. These could immediately be capitalised upon once the temporal HBI data became available.

One further, important unknown pertaining to HBI is the spatial scale across which within-mosquito population plasticity occurs. Over 50 years ago, Garrett-Jones described differing HBI estimates for mosquitoes collected from proximal locations [1]. Given the current concerns over altered biting behaviour potentially compromising recent gains in malaria burden reduction [21], a fuller comprehension of the scale and magnitude of this variability is timely. A recent study conducted in southern Ghana describes the successful piloting of a novel experimental design to address exactly this phenomenon [22]. It demonstrated that statistically significant alteration in host choice for *An. coluzzii* was detectable over a range of 250 metres [22]. Heterogeneity in mosquito biting rates has been demonstrated to be key to malaria transmission, first by theoretical work [23], but more recently with empirical studies using genotyping of blood-meals [24]. Future modelling frameworks will need to account for this additional form of village-level heterogeneity in biting behaviour.

To summarise, we presented the first systematic review of the HBI of the major African malaria vectors. We demonstrated that where mosquitoes are collected from (indoors versus outdoors) is significantly more associated with the HBI than which mosquito (sibling) species is collected. Some of the more important consequences to disease control of this behaviour are described. We further highlight some new theoretical and empirical developments that may improve both HBI assessment and how this metric can inform malaria control optimisation.

## Declarations

### Ethics approval and consent to participate

Not applicable

### Consent for publication

Not applicable

### Availability of data and material

All data generated or analysed during this study are included in this published article [and its supplementary information files].

### Authors’ contributions

LY conceived the study. JO, LFK and LY performed the systematic review and meta-analysis. All authors contributed to results interpretation and manuscript drafting.

## Acknowledgements

Not applicable

---

**Table S1:**
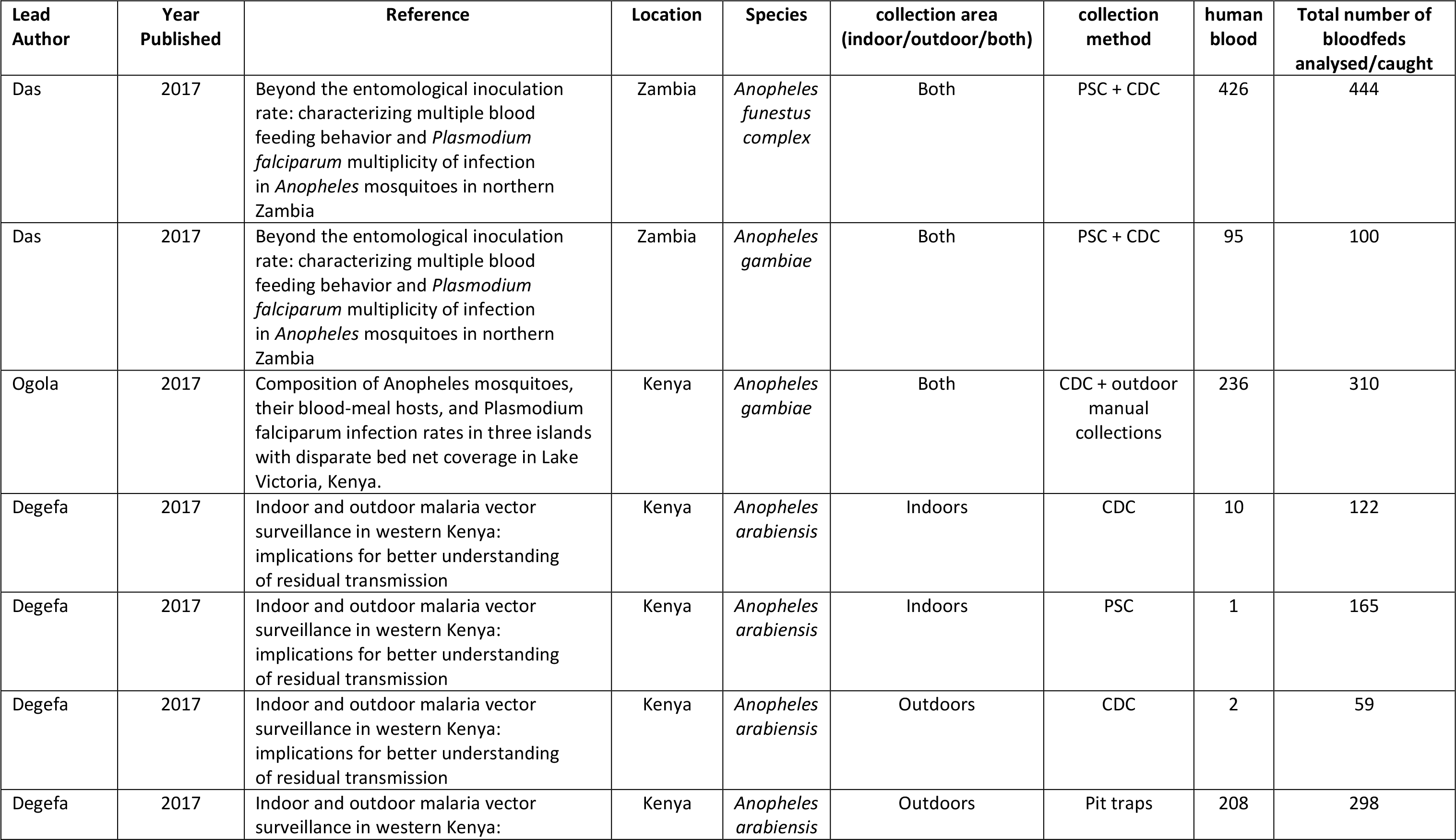
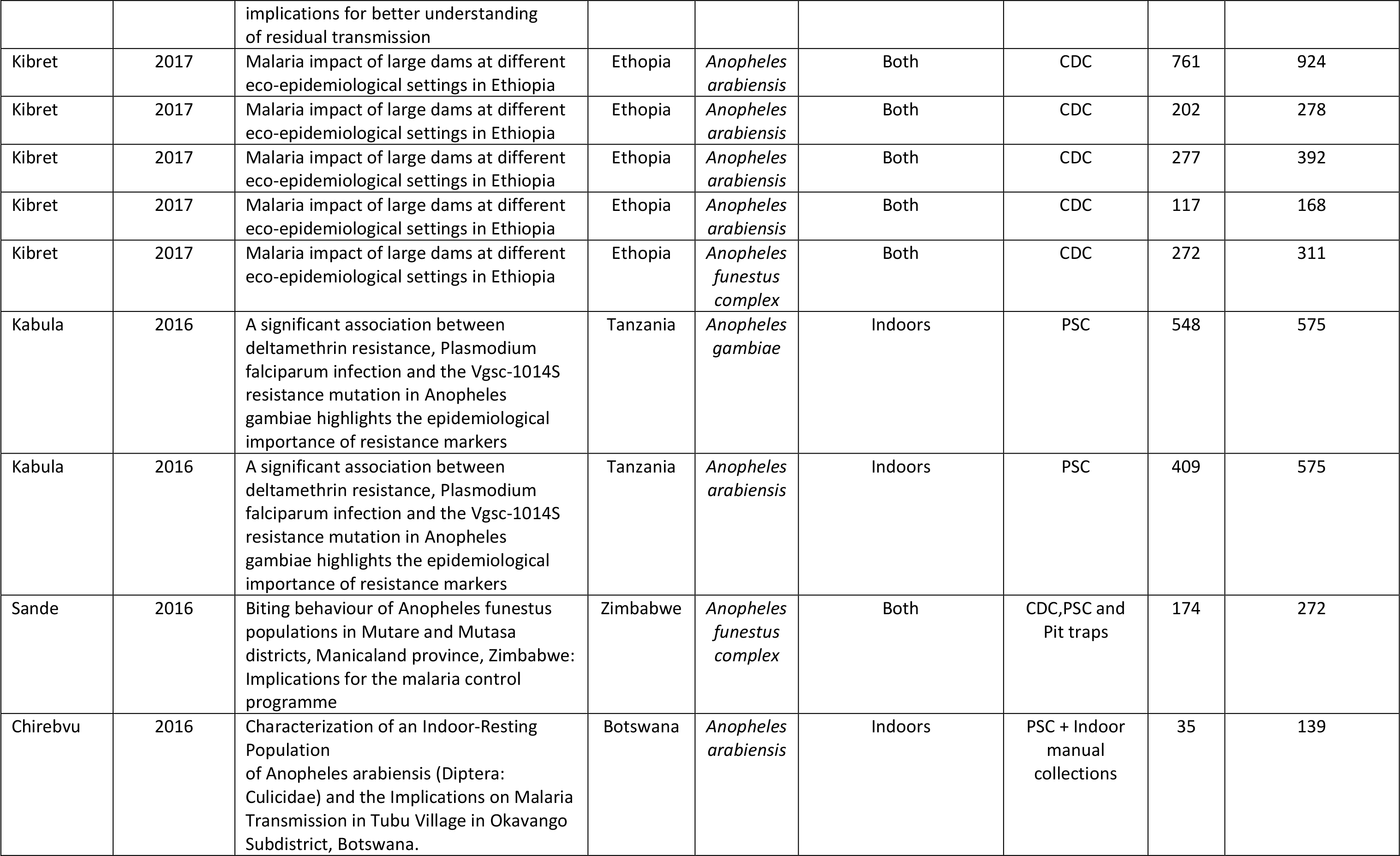
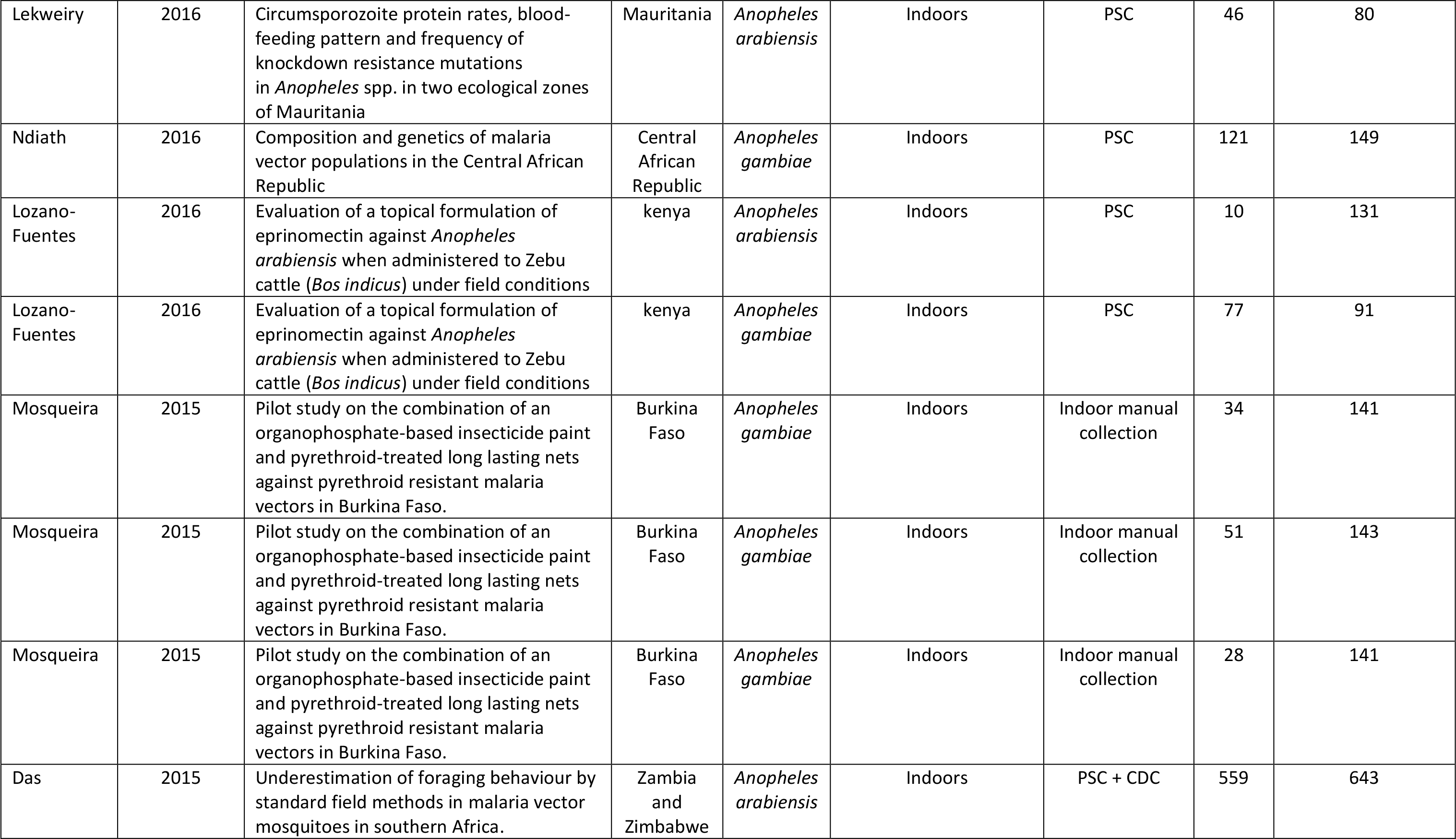
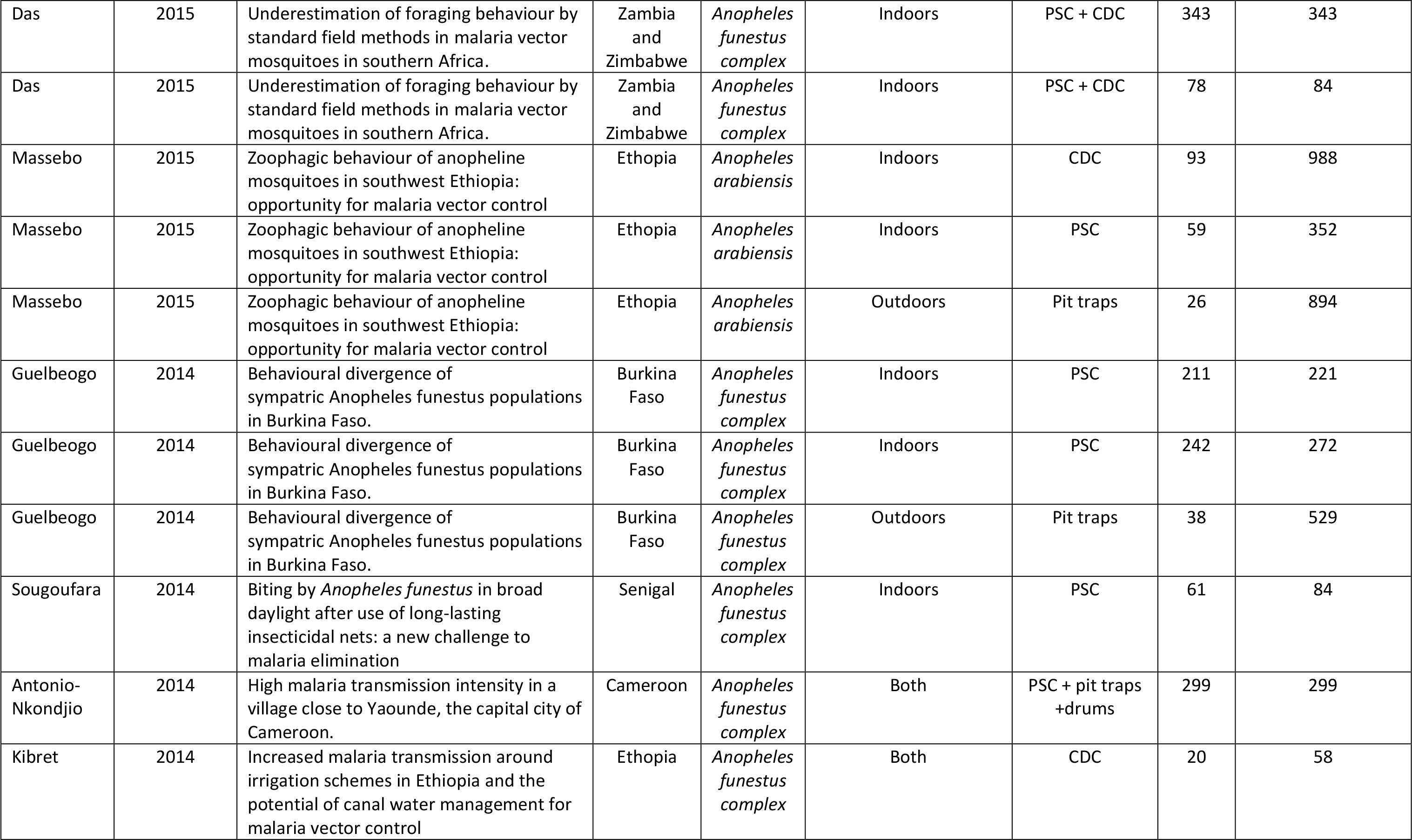
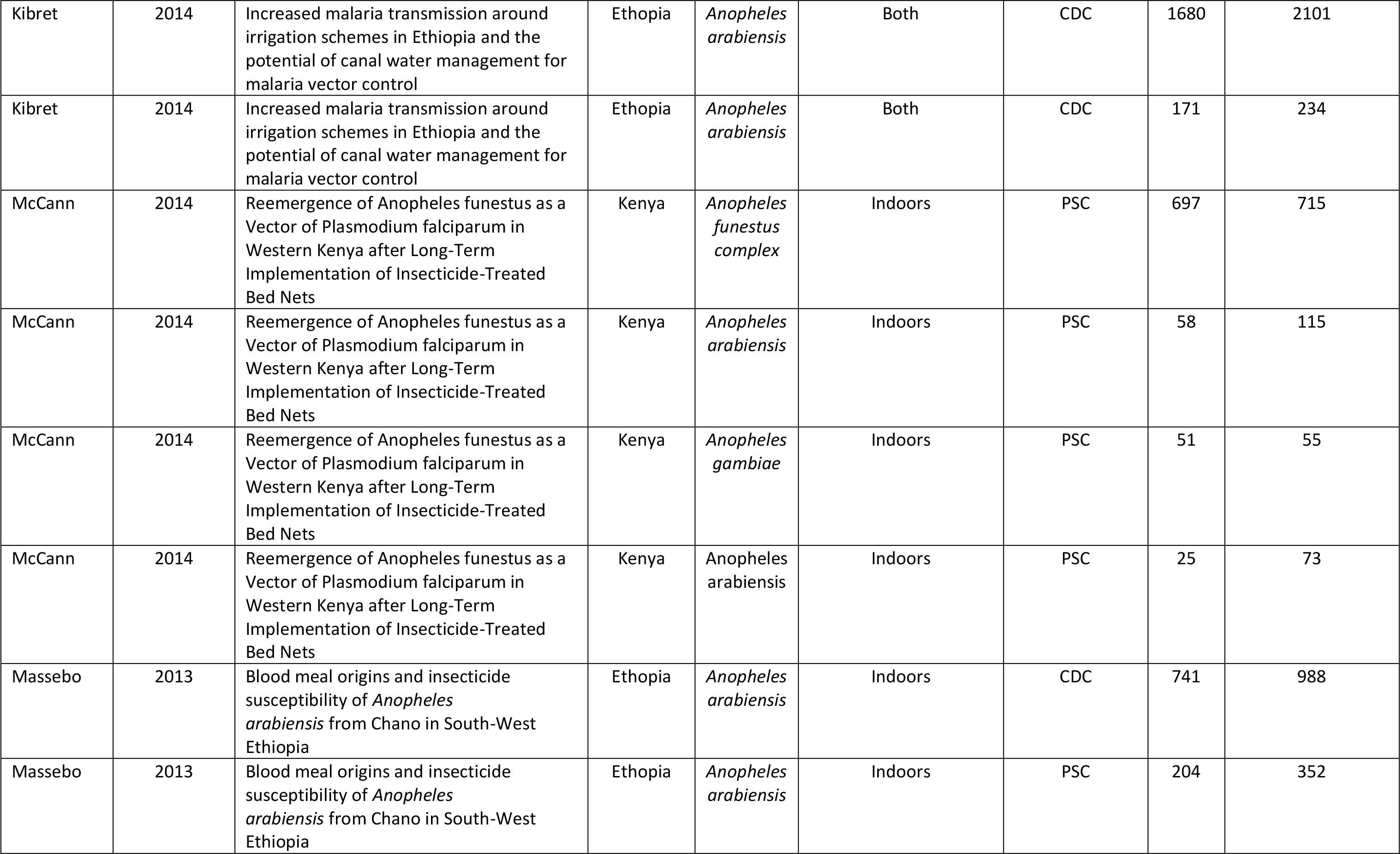
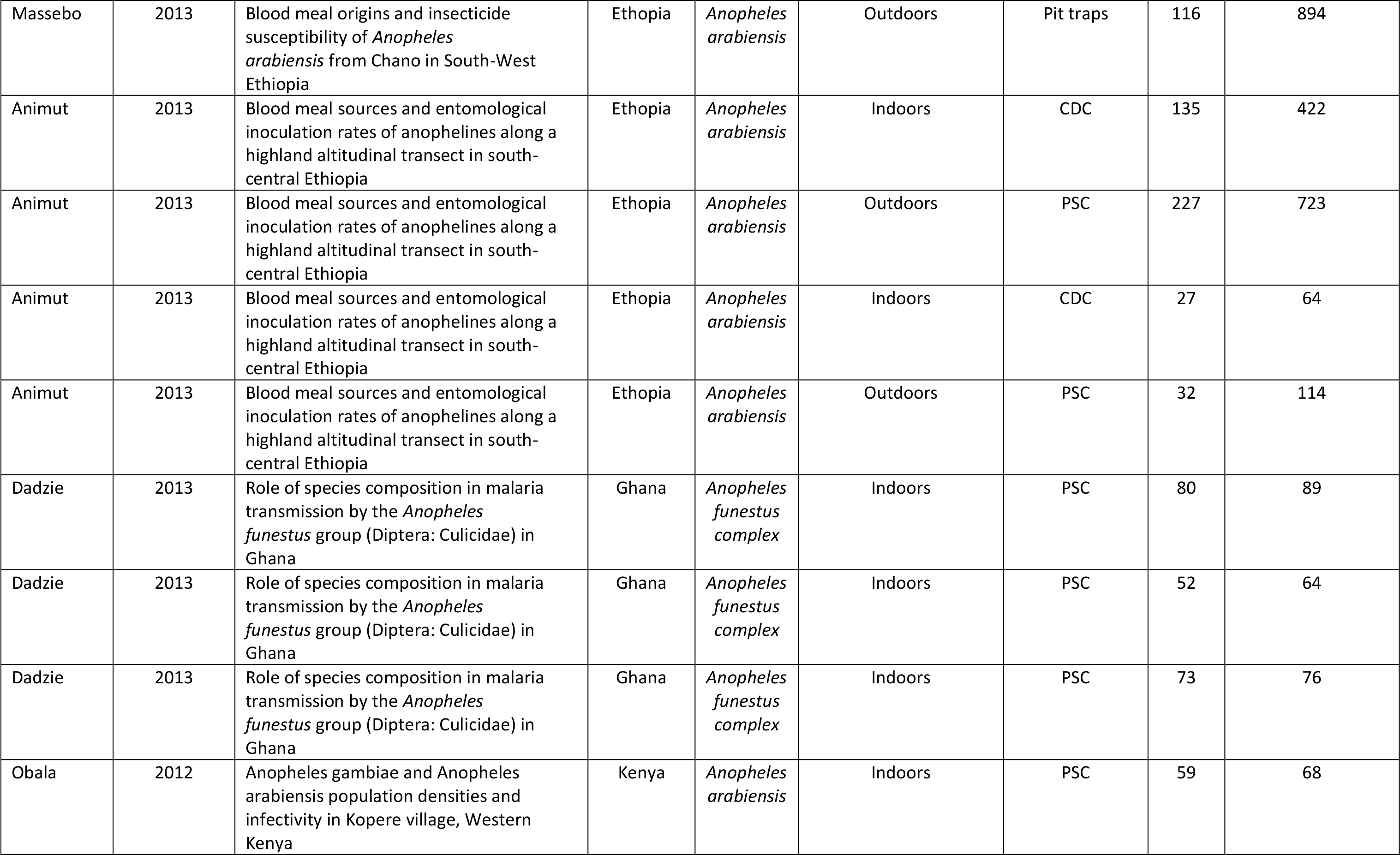
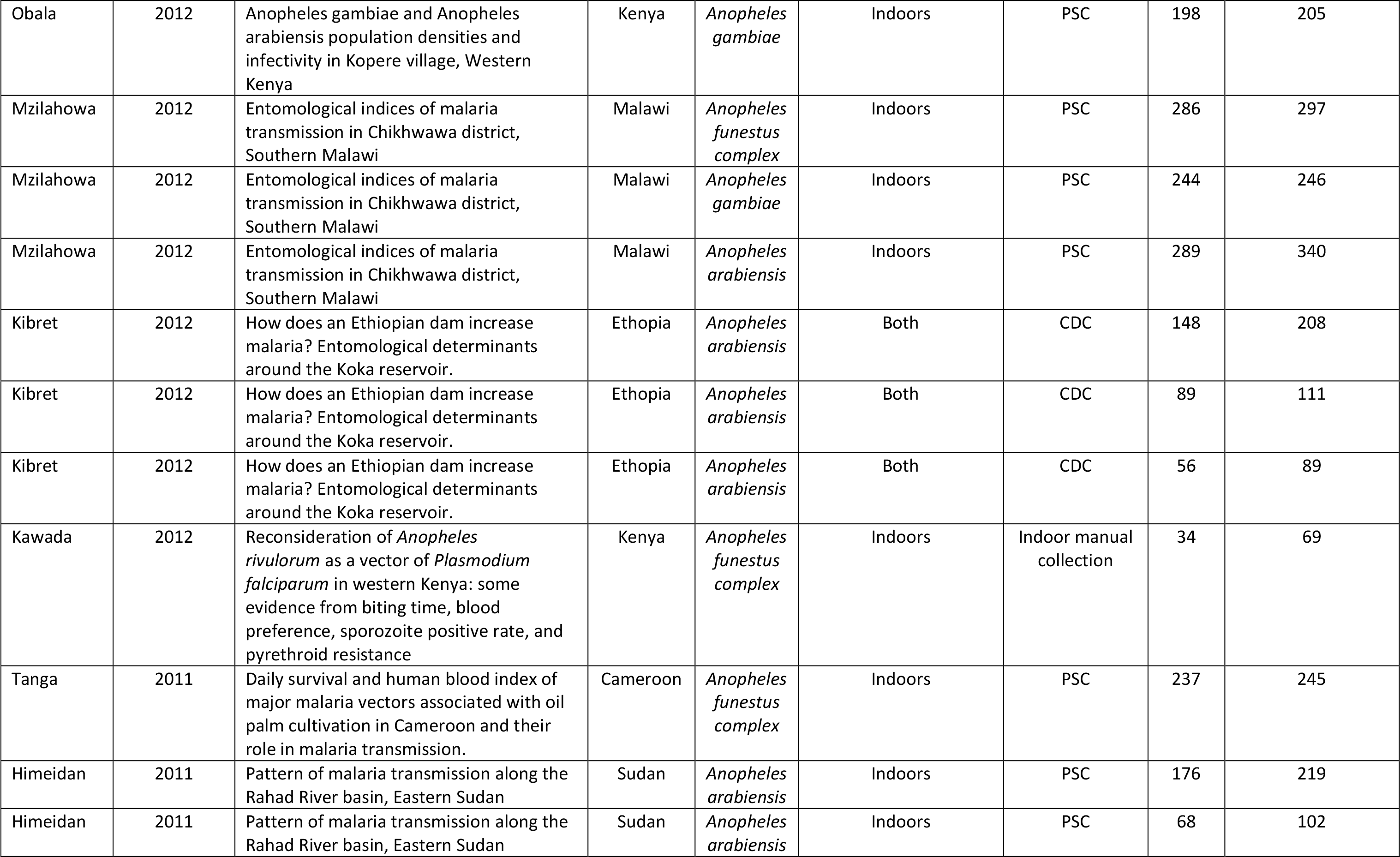
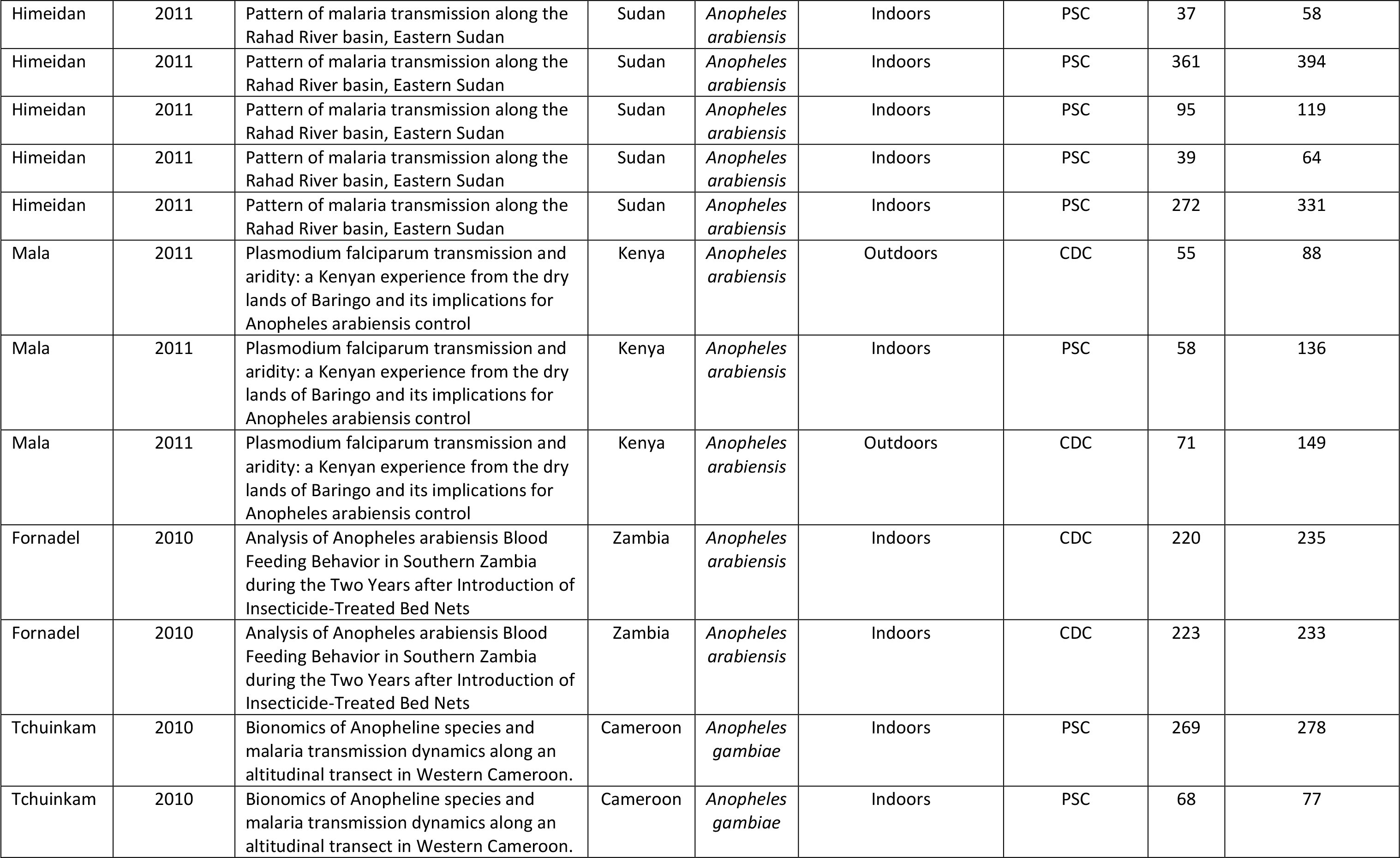
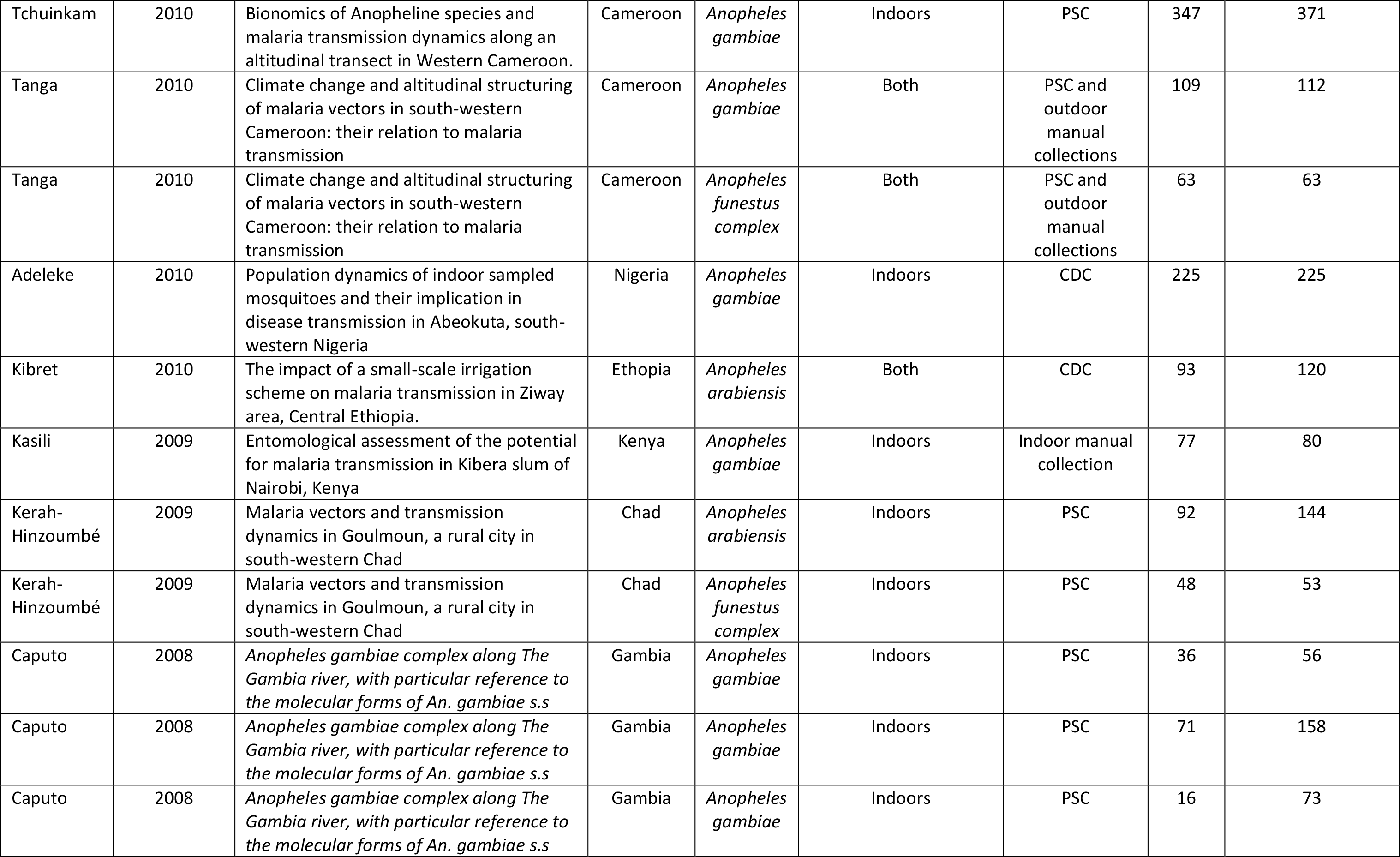
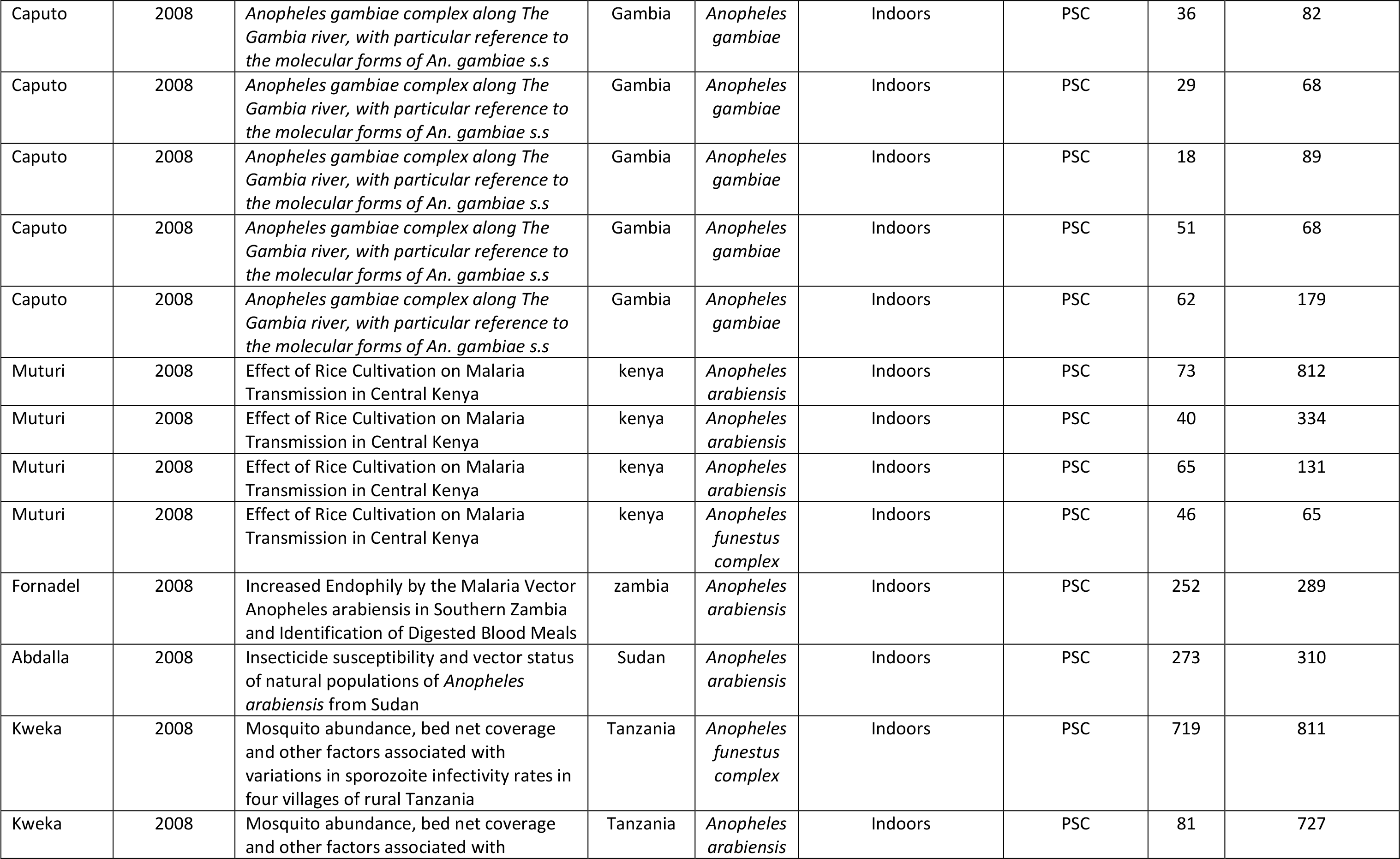
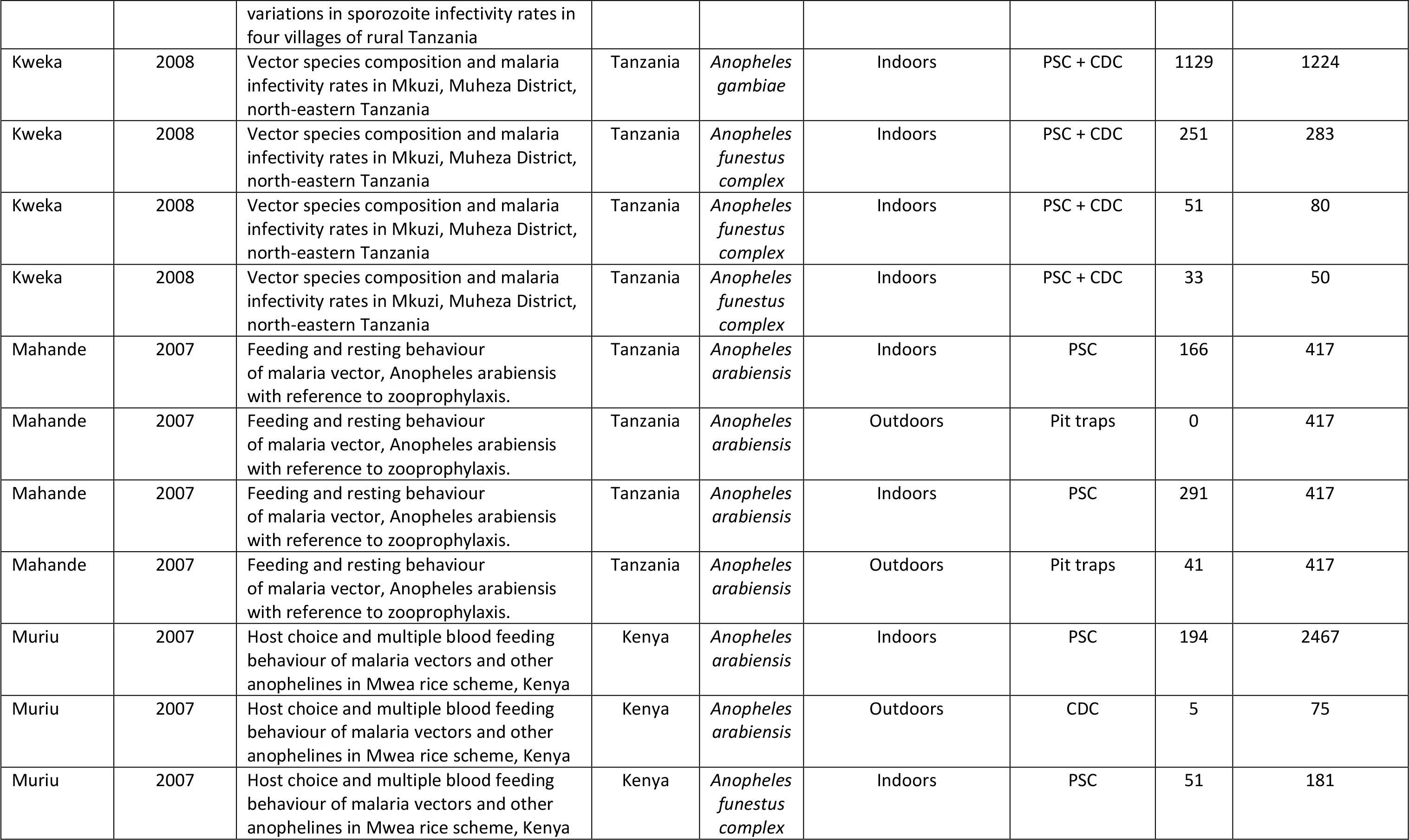
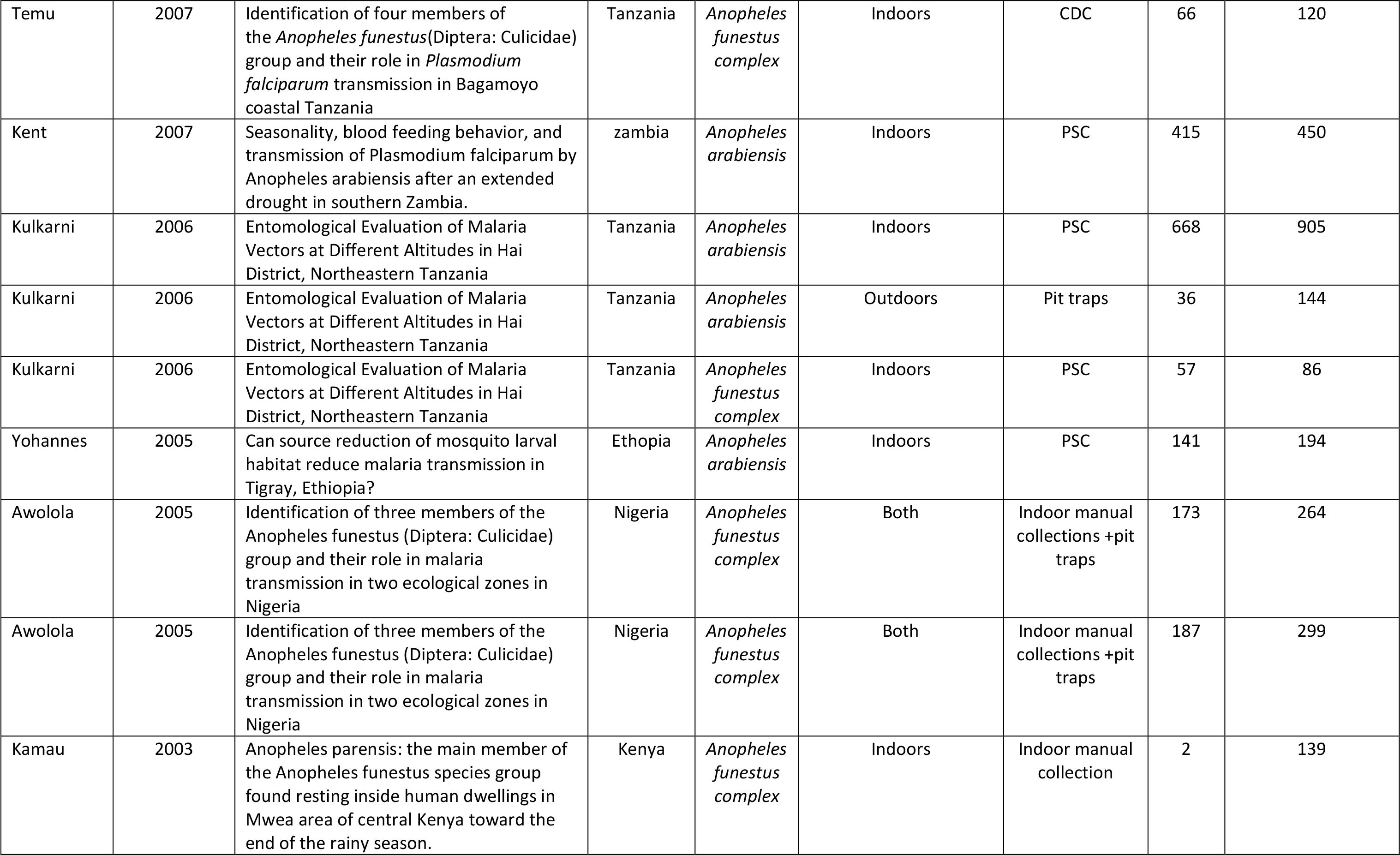
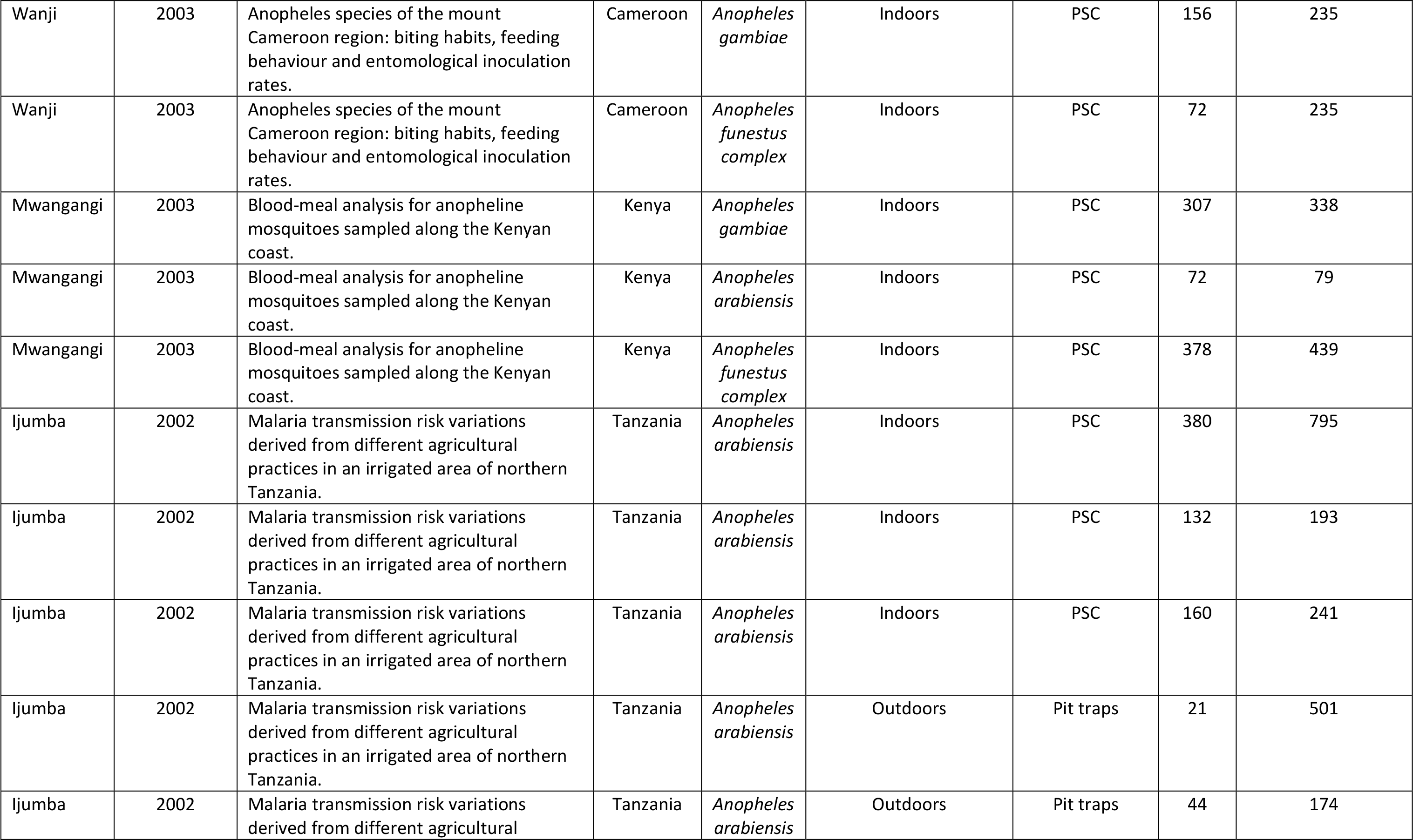
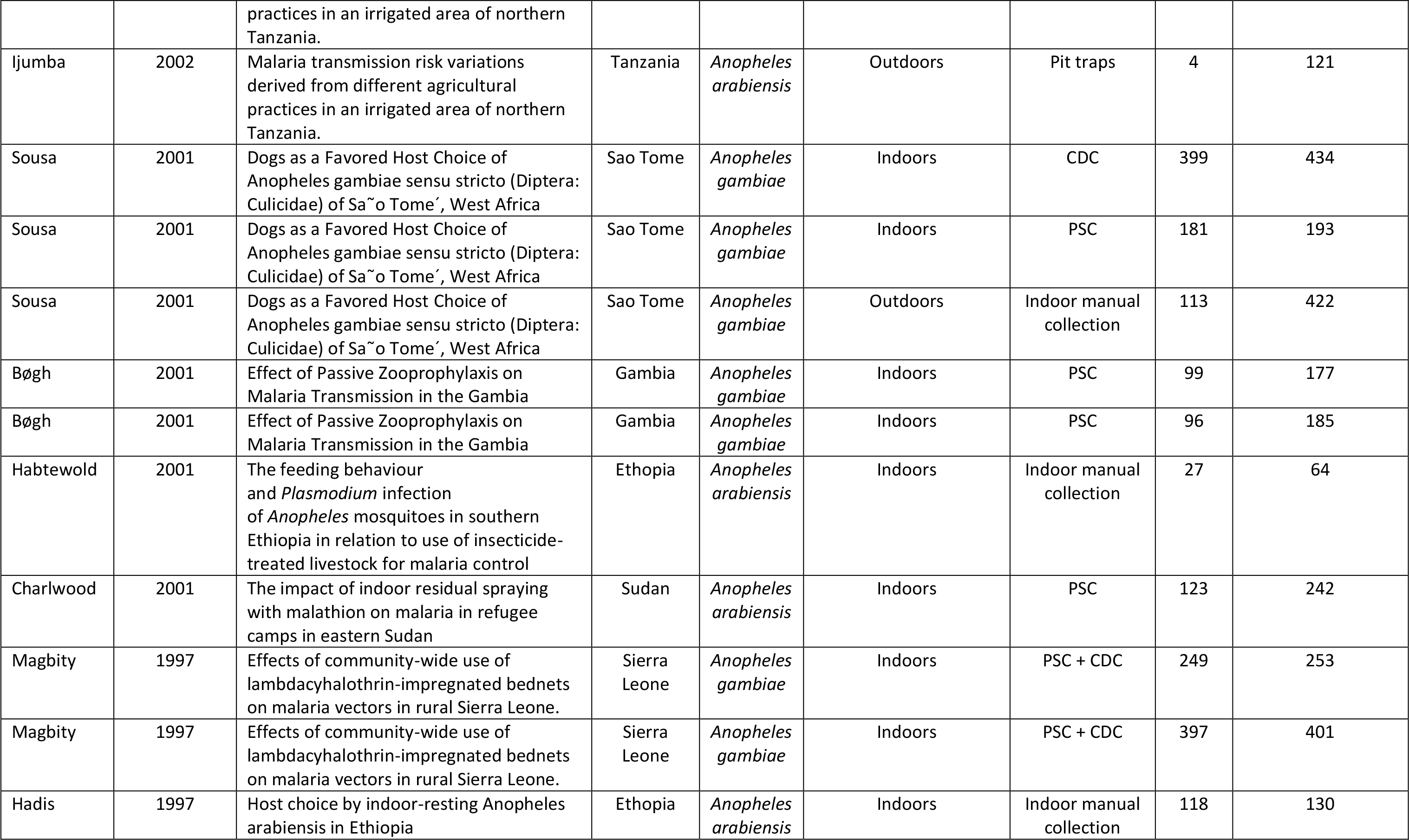
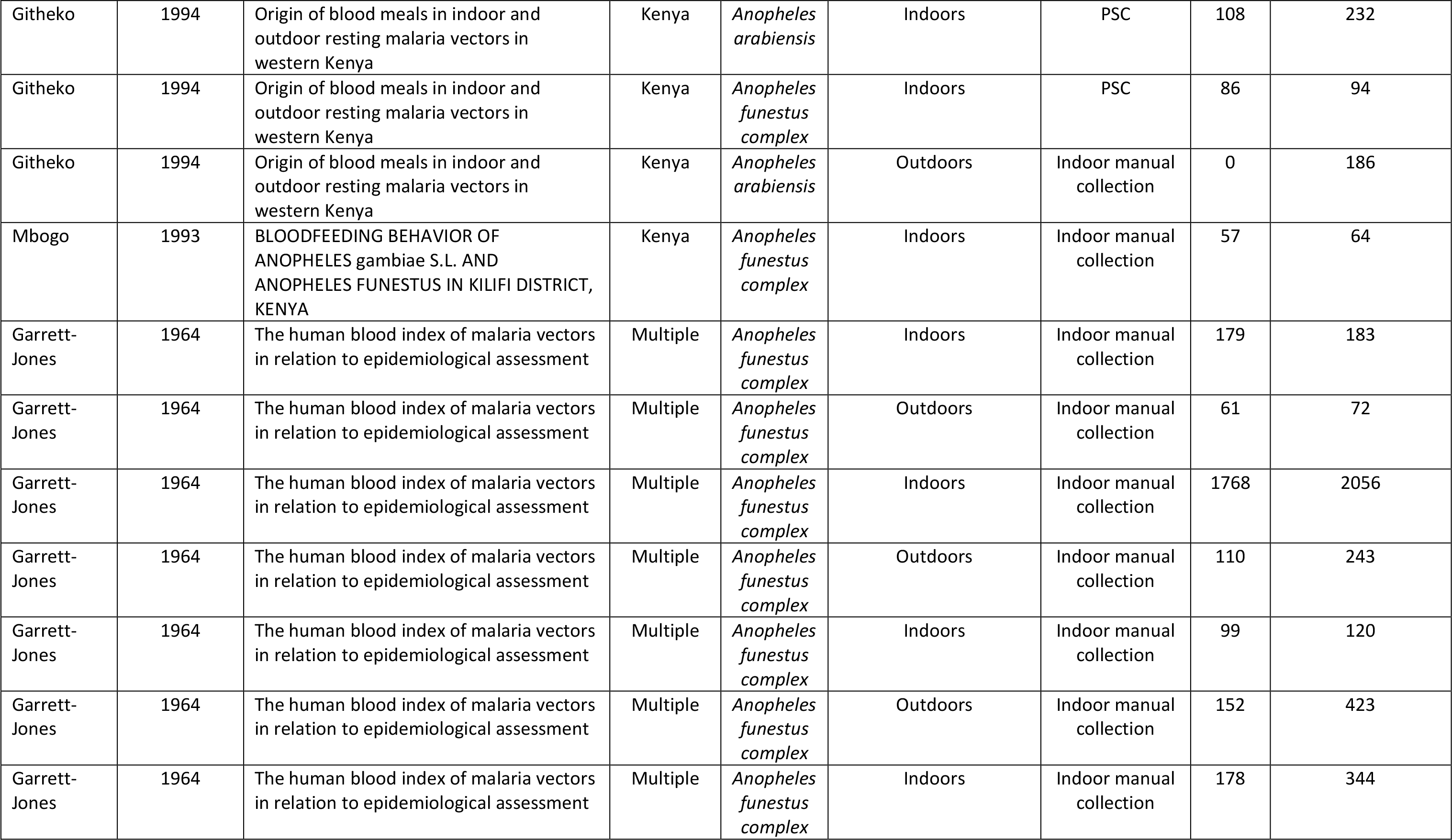
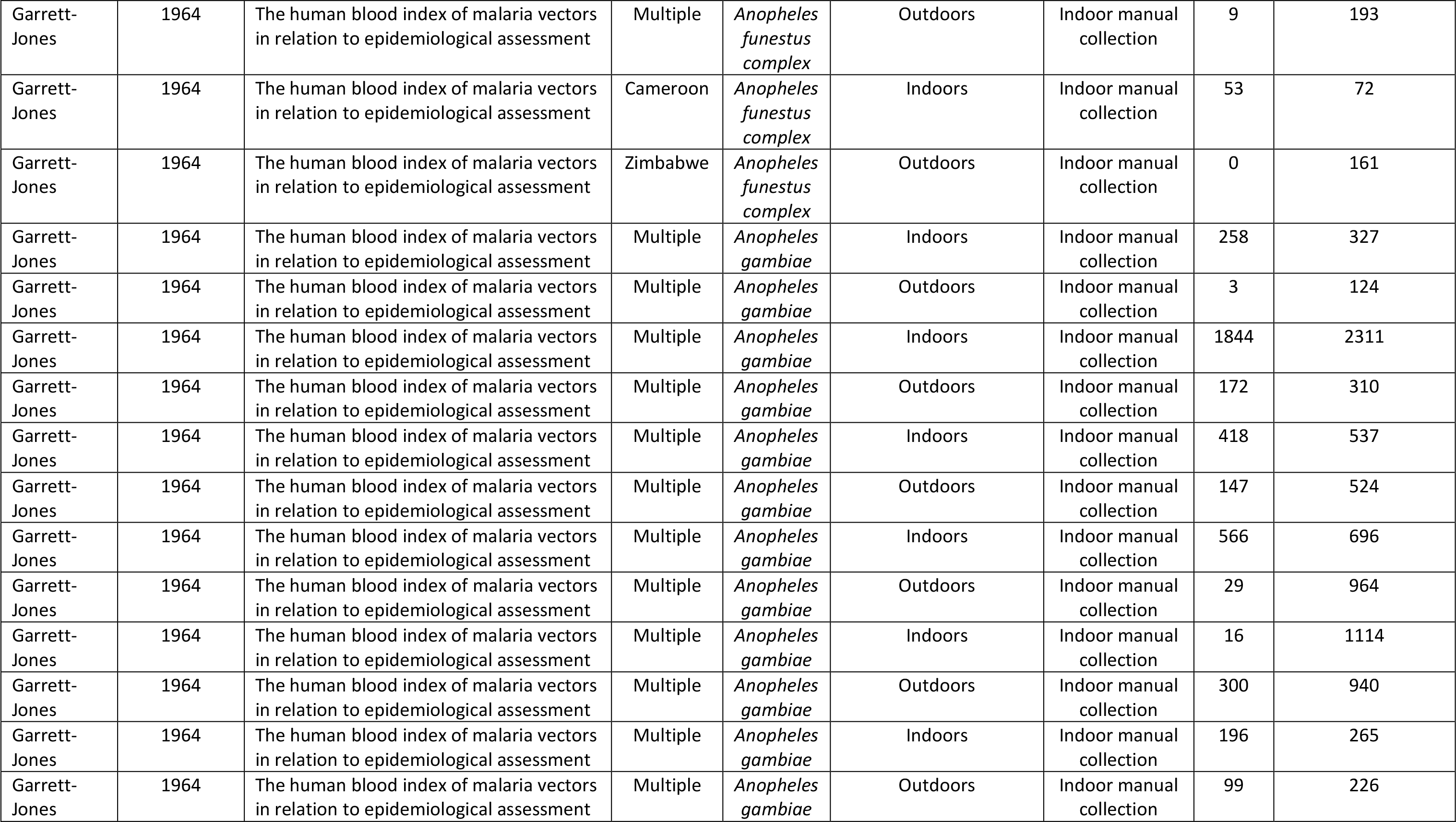
All eligible studies (and corresponding data points) retrieved from systematic search

